# *De novo* activated transcription of inserted foreign coding sequences is inheritable in the plant genome

**DOI:** 10.1101/2020.11.28.402032

**Authors:** Takayuki Hata, Naoto Takada, Chihiro Hayakawa, Mei Kazama, Tomohiro Uchikoba, Makoto Tachikawa, Mitsuhiro Matsuo, Soichirou Satoh, Junichi Obokata

## Abstract

The manner in which inserted foreign coding sequences become transcriptionally activated and fixed in the plant genome is poorly understood. To examine such processes of gene evolution, we performed an artificial evolutionary experiment in *Arabidopsis thaliana*. As a model of gene-birth events, we introduced a promoterless coding sequence of the firefly luciferase (*LUC*) gene and established 386 T2-generation transgenic lines. Among them, we determined the individual *LUC* insertion loci in 76 lines and found that one-third of them were transcribed *de novo* even in the intergenic or inherently unexpressed regions. In the transcribed lines, transcription-related chromatin marks were detected across the newly activated transcribed regions. These results agreed with our previous findings in *A. thaliana* cultured cells under a similar experimental scheme. A comparison of the results of the T2-plant and cultured cell experiments revealed that the *de novo*-activated transcription concomitant with local chromatin remodelling was inheritable. During one-generation inheritance, it seems likely that the transcription activities of the *LUC* inserts trapped by the endogenous genes/transcripts became stronger, while those of *de novo* transcription in the intergenic/untranscribed regions became weaker. These findings may offer a clue for the elucidation of the mechanism by which inserted foreign coding sequences become transcriptionally activated and fixed in the plant genome.

## Introduction

Genomes exhibit a steady state of the dynamic activity between the gain and loss of genes. Comparative functional genomics among closely related species revealed how genomes acquired such genetic novelty during their evolution, i.e., duplication–diversification, transposition, gene transfer or *de novo* origination [1–4]. Importantly, to become functional, new coding sequences must acquire promoters during their evolution. A gene promoter is a region in which transcription is initiated and is a central component of gene regulation [5,6]. In eukaryotes, promoter-specific sequences and chromatin marks are well characterized [5,6]. The study of the origin of genic promoters has been led by the examination of evolutionarily young genes. Notably, the *de novo* gene, whose coding sequence arises from an ancestral non-coding sequence, is a valuable model to study how newly originated coding sequences became transcribed [3,4,7,8]. *De novo* genes were often found from ancestral non-coding RNA genes [9,10], near the bidirectional promoters [9,11–14], and within enhancers [12,15–17], open-chromatin regions [16,17] and pervasively/divergent transcribed regions [18–25]. However, because the young genes found in present-day genomes became fixed thousands to millions of years ago, the molecular portraits of the newly originated promoters shortly after their birth are largely overlooked.

Differing from the comparative genomics approach, artificial evolutionary experiments can dissect the molecular mechanisms of how newly emerged coding sequences acquire their promoters in the timescales of molecular biology and genetics. Previously, to mimic a gene-birth event, we introduced exogenous promoterless coding sequences into *A. thaliana* T87 cultured cells and analysed their transcriptional fates at the genome-wide level [26] (unpublished data; available at https://doi.org/10.1101/2020.11.28.401992). Interestingly, we found that promoterless coding sequences became transcriptionally activated via two distinct mechanisms: (1) the so-called promoter trapping, in which integrated sequences capture the endogenous promoter activities of pre-existing genes/transcripts; or (2) *de novo* transcriptional activation, which occurs ubiquitously across the entire genome, and does so stochastically in about 30% of the integration events independently of the chromosomal locus [26]. The transcription start sites (TSSs) analysis revealed that the *de novo* transcription occurred from the 5’ close proximal region of the inserted foreign coding sequences [27]. We speculated that the insertion of exogenous coding sequences might activate local chromatin remodelling to shape the promoter-like chromatin configuration, resulting in such *de novo* transcriptional activation in the plant genome.

Could this *de novo* transcriptional activation be a causative mechanism by which newly originated coding sequences acquire their transcriptional competency in the plant genome evolution? Testing this possibility requires the assessment of the genetic behaviour of the *de novo* transcription over generations. The cultured cell-based experiment is not suitable for such scope because the cultured cells continue only the vegetative growth. In this respect, artificial evolutionary experiments with whole plants could provide clues to the above question. The plant body develops with the continuous formation of various tissues and organs from stem cells. Heterogeneity of the transcriptome and epigenome among these different tissues and developmental stages are well characterized in *A. thaliana* plants [28–31]. During plant reproductive development, dynamic chromatin remodelling including the localizations of DNA methylation and specific histone species occurs [32–36]. It is unpredictable from the cultured cell-based experiment how such chromatin remodelling could have an influence on the *de novo* transcription in the plant genome over generations.

In this study, we aimed to establish a model system to elucidate the mechanism by which inserted foreign coding sequences acquire their promoters and become fixed as functional genes in the plant genome. We carried out a large-scale promoter-trap screening in the T2 generation of *A. thaliana* plants under an experimental scheme similar to that used in our previous study of cultured cells [26]. By comparing the results obtained in plants with those of cultured cells, we concluded that *de novo* transcriptional activation together with chromatin remodelling is an inheritable phenomenon in the plant genome. After one generation, the transcriptional activities of introduced coding sequences trapped by endogenous genes/transcripts became much stronger, while those of the intergenic/untranscribed regions became much weaker. These findings may contribute to the elucidation of how newly emerged coding sequences become transcriptionally activated and fixed in the plant genome at their early evolutionary stages.

## Results

### Establishment of transgenic lines for large-scale promoter-trap screening in *A. thaliana*

To investigate the mechanism of promoter birth and their genetic behaviours beyond one generation, we performed a promoter-trap screening using *A. thaliana* plants under conditions that were essentially the same as those used in a previous study of cultured cells [26]. Based on *Agrobacterium*-mediated transformation [37], we introduced the promoterless coding sequence of a firefly luciferase (*LUC*) gene into *A. thaliana* (Fig 1). Each *LUC* gene was tagged by distinct short random sequences called ‘barcodes’ (Fig 1), which were used as identification codes for individual transgenic lines in the subsequent *in silico* analysis. In this study, to analyse the transgenic lines without the selection bias caused by *LUC* gene function, we screened the T1 seeds against the kanamycin (Km) resistance of the co-transformed expression cassette, rather than the strength of the LUC luminescence (Fig 1). Finally, we established a T2 generation of 386 individual transgenic lines (termed T2-plants hereafter).

**Fig 1.**
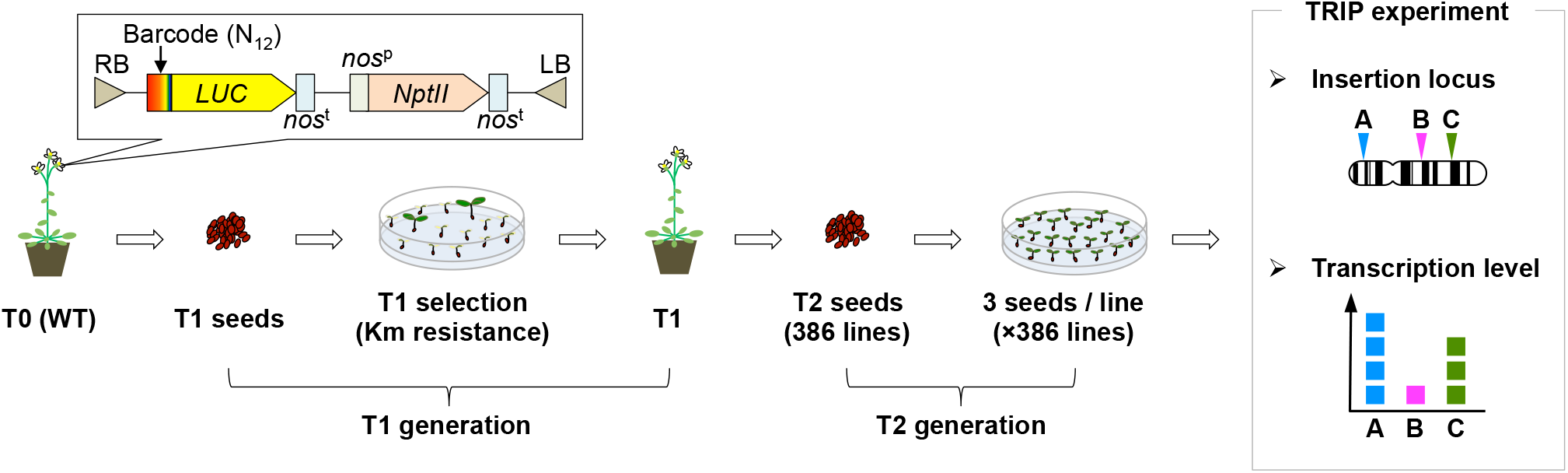
Experimental design of the promoter-trap experiment in *A. thaliana* plants. Schematic illustration of the TRIP experiment performed in the T2 generation of *A. thaliana* transgenic lines. T-DNA including a barcode, a promoterless *LUC* gene and an expression cassette with a Km-resistance gene was introduced into *A. thaliana* via *Agrobacterium*-mediated transformation. T2 seeds were harvested from Km-resistant T1 lines. Three seeds per T2 transgenic line were grown under the non-selective condition and subjected to subsequent locus and transcription-level analysis based on the TRIP method. *NptII*, neomycin phosphotransferase II; *nos*^p^, nopaline synthase promoter; *nos*^t^, nopaline synthase terminator.

### Genetic behaviours of *de novo*-activated transcription in *A. thaliana*

To identify the insertion loci and corresponding transcription levels of the individual *LUC* genes, we performed a massively parallel reporter assay based on the TRIP method [38].

First, three seeds per individual T2-plant were grown using the non-selective condition and seedlings were harvested as a mixed sample (Fig 1). Because the T2 generation is not homozygous, theoretically, one-fourth of T2 seeds were expected to be wild type (WT). However, as we grew three seedlings per line, no less than 98% of T2-plants (380/386) were expected to be recovered. In the TRIP method, individual transgenic lines are identified via *in silico* analysis based on the tagged barcode sequence of the reporter construct, as a molecular identifier. Specifically, we extracted DNA and RNA from the mixed samples and prepared the next-generation sequencing (NGS) library. For the determination of the insertion locus of each promoterless *LUC* gene, we performed inverse PCR followed by NGS, to read out the *LUC*– genome junction and barcode sequence. The relative transcription level of each *LUC* gene was determined utilizing NGS by counting the molecular abundance of each barcode in the RNA sample. Finally, each *LUC* gene-insertion locus and transcription level was assigned according to its barcode sequences. Note that T-DNAs are often inserted tandemly or with a large deletion on the reporter gene [39]. We carefully omitted such lines from further analysis because we could not determine their insertion loci uniquely. Based on this scheme, we determined individual insertion loci and corresponding transcription levels in 76 T2-plants (Figs 1 and 2A, and S1 Table). To confirm the results of the *in silico* analysis, we verified individual barcode sequences and insertion loci in randomly chosen T2-plants using Sanger sequencing and locus-specific PCR (S1 Fig). As shown in Fig 2A, promoterless *LUC* genes were inserted throughout the *A. thaliana* genome with low frequency in pericentromeric regions, which agreed with the reported preference of *Agrobacterium* T-DNA integration [40]. One-third of the 76 *LUC* genes (n = 27) were transcribed (Fig 2B). To examine further how these promoterless *LUC* genes became transcribed, we classified them according to their insertion types: an endogenous genic region with the sense (Genic Sense) and antisense (Genic AS) orientation, and the remaining intergenic regions (Intergenic). Based on this classification, the Genic Sense, Genic AS, and Intergenic types accounted for 26.3%, 21.1%, and 52.6% of the transcribed *LUC* genes, respectively (Fig 2C). Because the genic and intergenic regions of the *A. thaliana* genome have almost the same length [41], these results suggest that our established T2-plants exhibited no insertion-locus preference.

**Fig 2.**
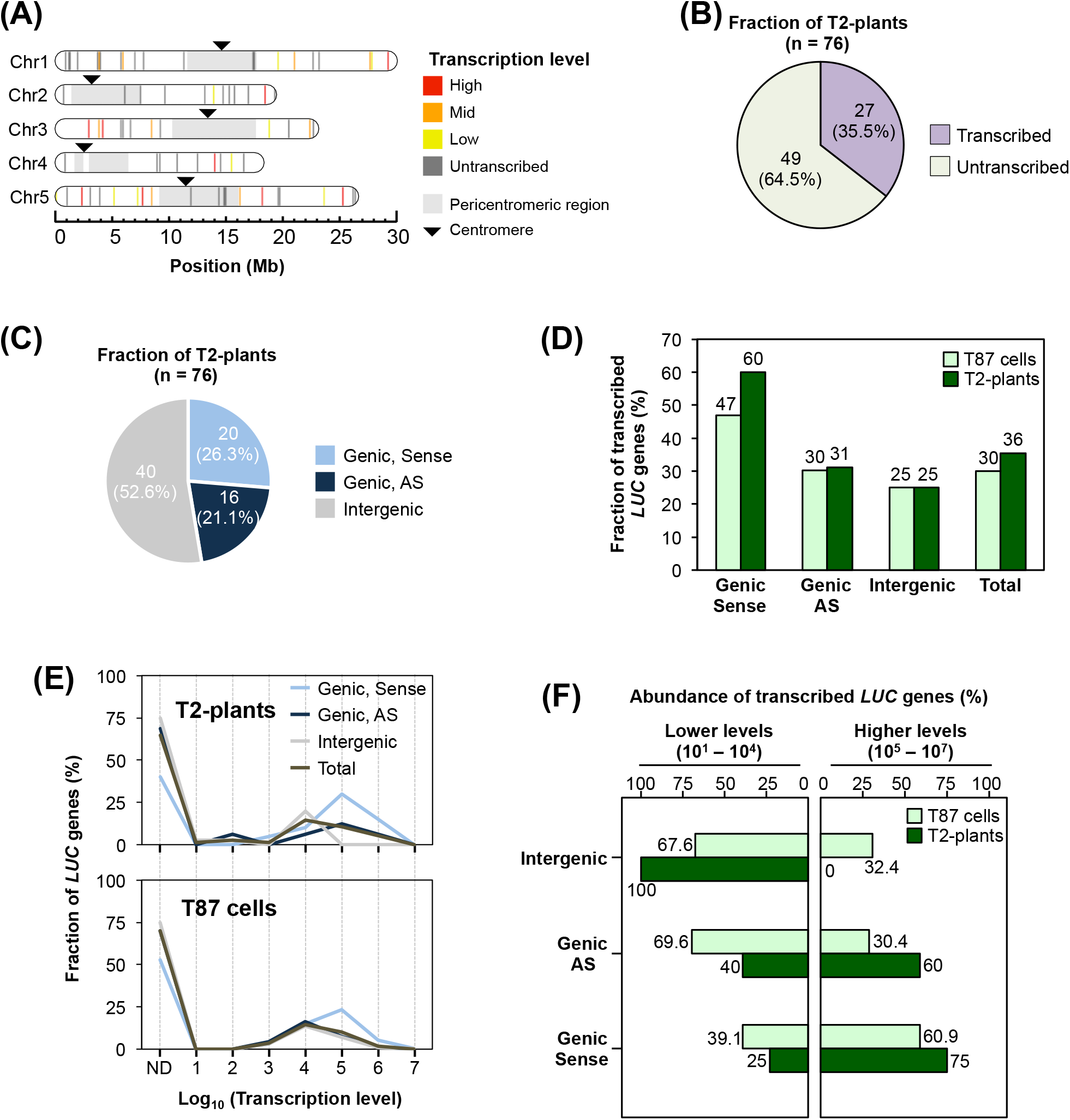
An artificial evolutionary experiment revealed the genetic behaviours of the activated transcription of coding sequences inserted in *A. thaliana* plants. **(A)** The insertion loci and transcription levels of determined T2-plants (n = 76) were mapped on the *A. thaliana* chromosomes. The coloured bars indicate individual insertion sites and corresponding transcription levels based on their percentiles (High: 100–67, Mid: 66–34, and Low: 33–1). (**B**) Classification of T2-plants according to their transcription. (**C**) Number of T2-plants according to their insertion types: Genic sense, AS (antisense) or intergenic. The definition of each type is provided in Materials and Methods. (**D**) Fraction of transcribed *LUC* genes among T2-plants (n = 76) and T87 cultured cells (n = 4,443) (Satoh *et al*., 2020) against each insertion type, as in (C). (**E**) Fraction of *LUC* genes in T2-plants (upper panel) and T87 cells (lower panel) (Satoh *et al*., 2020) against their transcription levels, as normalized using the total number of each insertion type as 100%. ND, untranscribed *LUC* genes. (**F**) The abundance of transcribed *LUC* genes in each insertion type was classified according to their transcription levels; lower (10^1^–10^4^) and higher (10^5^–10^7^), as in (E). Each frequency was normalized to the number of transcribed *LUC* genes in each insertion type, which was set as 100%.

Previously, based on the cultured cell experiment, we found that exogenously inserted promoterless genes became transcriptionally activated in two distinct types: promoter trapping and *de novo* transcriptional activation [26,27]. To examine whether similar transcriptional activation mechanisms occurred in our T2-plants, the abundance of the transcribed fraction was compared between the corresponding insertion types of T2-plants and cultured cells (Fig 2D and E). As shown in Fig 2D, ∽30% of the promoterless *LUC* genes were transcribed similarly in T2-plants and cultured cells (Fig 2D). Their relative transcription levels ranged from 10^1^ to 10^7^ orders, with a peak at 10^4^ (Fig 2E). Regarding the three insertion types, the abundances of transcribed *LUC* genes were almost the same in T2-plants and cultured cells, except for the Genic Sense type, in which the transcribed fraction was much greater in the T2-plants (Fig 2D). In both T2-plants and cultured cells, the Genic Sense type showed the highest transcriptional activity among the three insertion types, with 10^5^ as a peak (Fig 2E).

To highlight the differences between the cultured cells and T2-plants, we divided the transcribed *LUC* lines into two fractions: that with a lower transcription level (10^1^–10^4^) and that with a higher transcription level (10^5^–10^7^). As shown in Fig 2F, the relative abundances of the higher and lower fractions in T2-plants exhibited a greater bipolarization than they did in cultured cells; *LUC* transcription became much stronger in the Genic Sense and AS types, while it became much weaker in the Intergenic type (Fig 2F). As the Genic Sense and AS types were transcribed presumably by trapping the transcriptional activities of endogenous genes [27], these features suggested that gene-trapping events are more prone to occur in plants than in cultured cells. Conversely, a type of transcriptional repression might have occurred on the Intergenic type in the T2 generation.

Taken together, these results suggest that the transcriptional behaviours of the promoterless *LUC* genes are remarkably similar between the T2-plants and the vegetatively growing cultured cells (Fig 2D and E). Therefore, it is likely that *de novo* transcriptional activation events are not specific to the vegetatively growing cultured cells; rather, they seem to be an inheritable phenomenon through a plant’s generation.

### Comparison of *LUC* transcription with inherent transcriptional status

Are there any other similarities/differences between T2-plants and cultured cells? To address this question, we next focused on the correlation of the transcriptional status between the *LUC* genes and the corresponding WT loci. For this, we prepared a dataset of the transcribed regions of the WT genome using the publicly available RNA-seq data of *A. thaliana*, which were obtained using growth conditions similar to those used here. The WT dataset represents mostly (97.8%) the annotated genic regions, which cover 70.4% (19,308/27,416) of the annotated protein-coding genes. We classified the *LUC* insertion loci into four types by the combination of the transcriptional status of the *LUC* genes and the corresponding WT loci: (i) a *LUC* gene was transcribed in the WT transcribed region; (ii) a *LUC* gene was untranscribed in the WT transcribed region; (iii) a *LUC* gene was transcribed in the WT untranscribed region; and (iv) a *LUC* gene was untranscribed in the WT untranscribed region. The relative abundance of each type in T2-plants and T87 cells [26] was as follows: (i) 14.5% and 7.8%, (ii) 7.9% and 8.2%, (iii) 21.1% and 22.3% and (iv) 56.6% and 61.7%, respectively (Fig 3A). Based on these data, we evaluated the transcriptional activation rates in the WT untranscribed regions more precisely. We then redrew Figure 3A using the sum of types (iii) and (iv) as 100% (Fig 3B). In this data presentation, the transcriptional activation frequency was surprisingly similar between T2-plants and cultured cells (27.1 vs. 26.5 in Fig 3B), which supports the contention that *de novo* transcriptional activation in the untranscribed region occurs similarly in plants and cultured cells. Next, we performed the same analysis for the WT transcribed regions (types (i) and (ii)), and showed that the transcriptional activation frequency was higher in the T2-plants than in the cultured cells (66.0 vs. 48.7 in Fig 3C). This feature of the transcribed regions (Fig 3C) was reminiscent of the feature of the annotated genes (Fig 2D), because most of the WT transcribed regions (97.8%) represent annotated protein-coding genes. They both showed that the *LUC* inserts in the genic regions were activated more frequently in the T2-plants than in the cultured cells. The possible explanations for this feature from the viewpoint of plant life cycles and Km-based selection during the T2-plants establishment are referred to in the discussion section.

**Fig 3.**
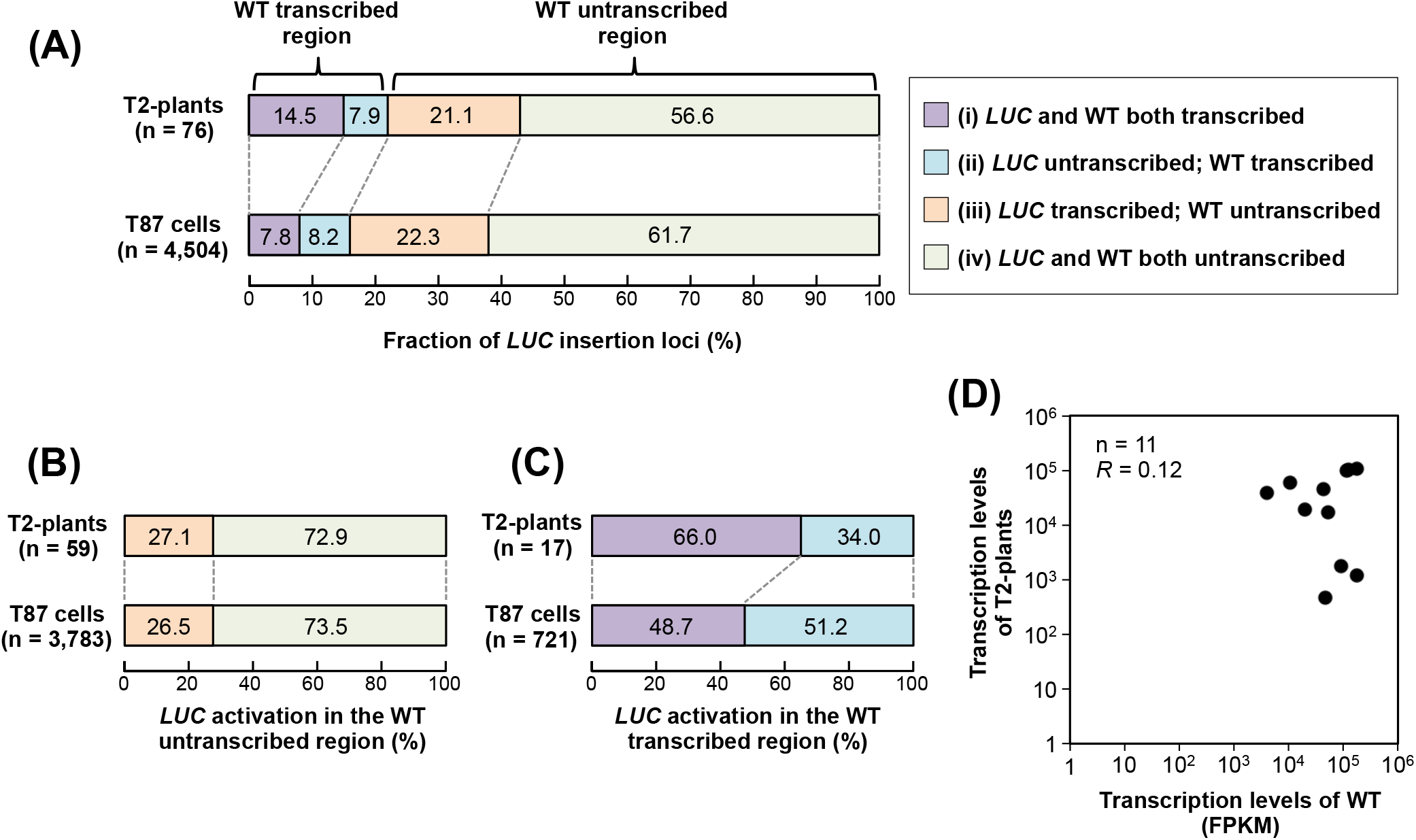
Transcription status of *LUC* insertion loci in T2-plants and WT plants. **(A)** *LUC* insertion loci were classified by the combination of the transcription status of the *LUC* gene and the corresponding WT locus as follows; (i) *LUC* gene and the corresponding WT locus were both transcribed; (ii) *LUC* gene was untranscribed, but WT locus was transcribed; (iii) *LUC* gene was transcribed, but WT locus was untranscribed; and (iv) *LUC* gene and WT locus were both untranscribed. Upper panel, T2-plants; lower panel, T87 cells (Satoh *et al*., 2020). (**B**) Relative fractions of types (iii) and (iv) normalized to the sum of their abundances in (A), as 100%. (**C**) Relative fractions of types (i) and (ii) normalized to the sum of their abundances in (A), as 100%. (**D**) The transcription levels of T2-plants and corresponding inherent transcripts in WT plants were compared. *R*, Spearman’s correlation test.

Generally, in promoter-trapping experiments, the expressed reporter genes are expected to reflect the activities of trapped endogenous promoters [42]. However, we previously found that the transcription levels of promoterless *LUC* genes did not reflect those of their inherent endogenous transcripts in the experiment that used cultured cells [26]. To confirm whether this feature was specific to the vegetatively growing cultured cells, we compared the transcription levels between T2-plants and their corresponding regions in the WT genome. We found that there was no correlation between them (Fig 3D). Thus, the observation that the trapping type of newly activated transcription events did not retain their inherent transcriptional status, at least in our experimental conditions, appeared to be a general feature of the plants and cultured cells. As insertions of the fragments were likely to disrupt the transcriptional activities of given loci, this result suggests two possibilities: (1) the original transcriptional activities had not yet been recovered in the vegetative propagation or within one generation; or (2) the transcriptional activities were overwritten by the *de novo*-activated transcription.

### Chromatin remodelling occurred across the newly activated transcribed regions

Eukaryotic transcription is regulated by the control of the localization of transcription-related chromatin marks [5,6]. Therefore, next, we focused on the chromatin configuration around *LUC* inserts to examine whether the transcribed T2-plants were regulated by such chromatin marks. First, we screened T2-plants according to the following criteria: the existence of transcription evidence in the TRIP experiment, and an unlikeliness to be affected by the pre-existing promoters or transcription units. Based on these criteria, we finally selected two lines: T2:161 and T2:205 (Fig 4A and B). The T2:161 line was classified as a Genic AS type in which the *LUC* insert was found in the opposite strand of an endogenous gene (AT3G23750) (Fig 4A). In the T2:205 line, the *LUC* insert was located in an intergenic region, in which an endogenous gene (AT5G01110) was detected downstream of the *LUC* insert on the opposite strand (Fig 4B). Transcription of inserted promoterless *LUC* genes was verified in both lines by reverse-transcription quantitative-PCR (Fig 4C, and S2 Fig), whereas no RNA-sequencing reads were mapped on the same strand of each *LUC* insert in the corresponding WT genome, indicating that they were inherently untranscribed regions. For these two lines, we scanned the localization of chromatin marks around the *LUC* insertion loci and compared them with those obtained from the WT genome. In this study, we analysed three transcription-related chromatin marks: methylated cytosine (mC), lysine 36 tri-methylation of histone H3 (H3K36me3), and the histone variant H2A.Z. In the WT genome, enrichments of mC and H3K36me3 were observed within the gene bodies of AT3G23750 and AT5G01110, respectively (Fig 4D and E, upper and middle panels), which agreed with the general properties of these chromatin marks [43,44]. However, in the T2-plants, these two chromatin marks were not found within the *LUC* gene bodies (Fig 4D and E, upper and middle panels). Although weak signals were observed 200 bp upstream from the *LUC* insert in the T2:161 line (Fig 4D, upper and middle panels), they reflected the chromatin marks of the WT allele in the T2-plant, because these plants were not homozygous. Conversely, the localization patterns of the H2A.Z variant were clearly different from those of the other two chromatin marks (Fig 4D and E, lower panels). Both lines showed significant enrichments of H2A.Z throughout the *LUC* gene bodies, while there were almost no H2A.Z signals in the corresponding regions in the WT genome (Fig 4D and E, lower panels). Although H2A.Z is a marker histone for the promoter region, it also appears in the gene bodies of genes with low expression [45–47]. In addition, mC and H3K36me3 were reportedly deposited within a gene body in a transcription-coupled manner [48], which would be undetectable in the low-expressed genes [49]. Thus, these distribution patterns of chromatin marks in the T2-plants were plausible because the transcriptional strength of these two lines was low compared with that of the constitutive genes (Fig 4C, and S2 Fig).

**Fig 4.**
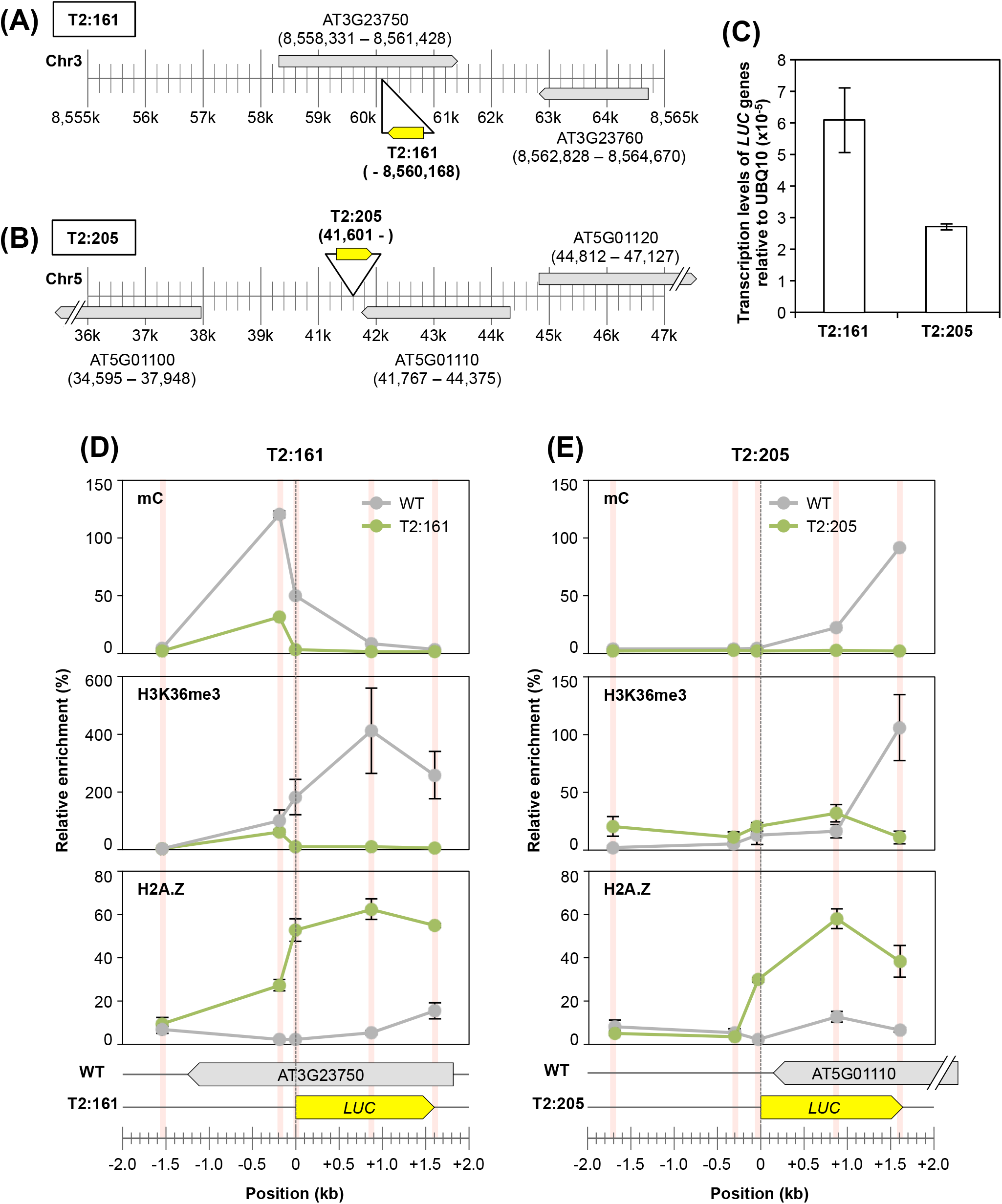
Localization analysis of chromatin marks in selected T2-plants. (**A** and **B**) Locus details of the (A) T2:161 and (B) T2:205 lines. The genomic loci of *LUC* inserts are represented as the individual position of the RB–genome junction. (**C**) Transcription levels of the T2:161 and T2:205 lines relative to the endogenous *UBQ10* gene (AT4G05320). (**D** and **E**) Localization patterns of three chromatin marks (mC: upper panel; H3K36me3: middle panel; and H2A.Z: lower panel) around individual *LUC* insertion loci of the (D) T2:161 and (E) T2:205 lines. Individual localization signals were normalized to the enrichment of the control locus of each chromatin mark (see Materials and Methods) as 100%. The red bars indicate the analysed positions, which were normalized to the genomic position of the start codon of *LUC* inserts as zero. Error bar, ±SD of two biological replicates.

In the T2:161 line, H2A.Z was newly localized 200 bp upstream from the *LUC* insert (Fig 4D, lower panel), which suggests that chromatin remodelling occurred even outside the *LUC* insert. We hypothesized that H2A.Z is localized throughout the transcribed region of the *LUC* insert. To confirm this hypothesis, next we analysed the TSS of *LUC* inserts. However, it was challenging to determine the TSSs of T2-plants using general methods [50,51] because of the low transcription levels of these plants. The template-switching method has the advantage of yielding full-length cDNAs from low-input RNA [52]. In this study, we applied inverse PCR to this template-switching method to specifically amplify the full-length cDNAs of *LUC* genes. Based on this method, we analysed TSS distribution in T2-plants. Unfortunately, the transcription level of the T2:205 line was too low to obtain any TSS signals. Conversely, in the T2:161 line, a TSS was found ∽1.1 kb upstream of the *LUC* insertion locus (Fig 5A, and S3 Fig). Sanger sequencing revealed that this transcript was spliced (Fig 5A, and S3 Fig). We reanalysed the distribution profiles of H3K36me3 and H2A.Z around the determined TSS (Fig 5B). There was no significant enrichment of H3K36me3 around the *LUC*-TSS, as the enrichment levels were almost the same among the transgenic plants and the WT genome (Fig 5B, upper panel). In contrast, we observed that H2A.Z was newly localized starting from the *LUC*-TSS, whereas H2A.Z was not observed in the corresponding locus in the WT genome (Fig 5B, lower panel).

**Fig 5.**
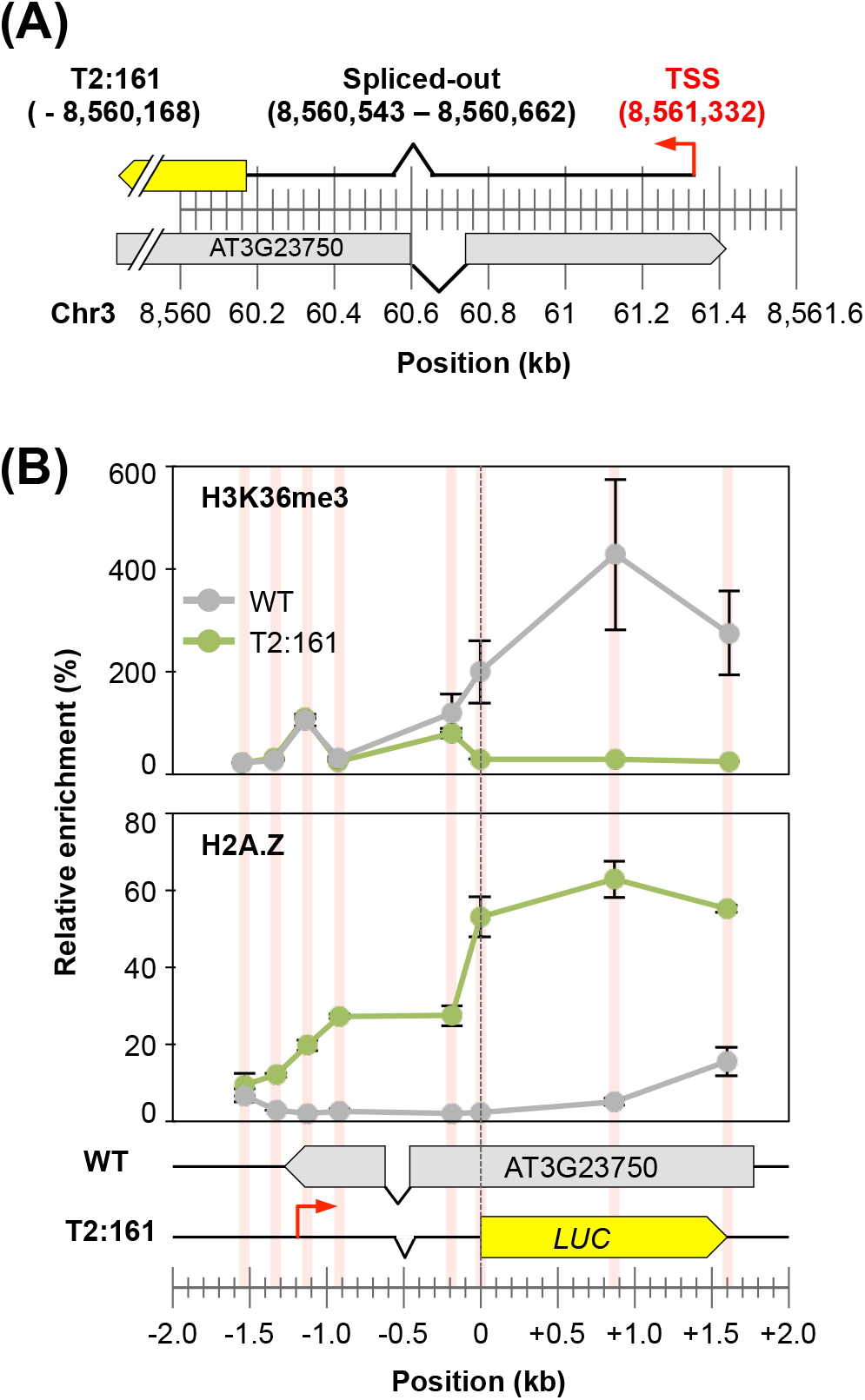
Localization of chromatin marks around the transcription start site of the T2:161 line. (**A**) Transcription start site of the T2:161 line, as determined using a template-switching-based method. (**B**) Localization analysis of H2A.Z and H3K36me3 in T2:161 and WT plants, as in Fig 4D.

Overall, the chromatin and TSS analyses revealed that exogenously inserted promoterless genes acquired a brand-new chromatin configuration, and that such chromatin remodelling occurred throughout the newly activated transcription unit. In addition, this chromatin remodelling might have been involved in the transcriptional behaviour of the trapping type of *LUC* transcription (Fig 3D): *de novo*-activated transcription events concomitant with the chromatin remodelling might overwrite their inherent transcriptional status.

## Discussion

In this study, based on the large-scale promoter-trap screening of *A. thaliana* plants, we demonstrated the genetic behaviour of the newly activated transcription of exogenous genes. A comparison with the results of a previous study using cultured cells [26] showed that *de novo* transcriptional activation is an inheritable phenomenon of the plant genome (Figs 1–3). We also demonstrated that chromatin remodelling occurred across the transcribed regions of the inserted coding sequences in the selected two transgenic lines (Figs 4 and 5), which probably regulated the newly activated transcription of these loci by overwriting the inherent chromatic and transcriptional status.

In the T2:161 line, the TSS was located on the 3’
s end of an endogenous gene (AT3G23750), where no detectable transcripts existed in the WT genome (Fig 5A). It is plausible to propose that this was caused by activating (rather than trapping) a cryptic antisense transcript of the given locus [27]. Conversely, we speculated that the T2:205 line may be transcribed from a *de novo*-activated TSS located in the proximal intergenic region, although we could not identify this TSS in this study. This speculation was based on a previous finding from the cultured cell experiment: *de novo* TSS occurs about 100 bp upstream of the inserted coding sequences in the intergenic region [27]. The localization pattern of H2A.Z in the T2:205 line agreed with this prediction, as the H2A.Z signal clearly dropped to almost zero at 200 bp upstream of the *LUC* insert (Fig 4E).

Generally, in promoter-trap screening, transgenic lines are screened based on the expression of the inserted promoterless reporter genes [42]. In contrast, we did not carry out the screening of T2-plants according to the expression of *LUC* genes; rather, we selected them according to the activity of the co-transformed Km-resistance gene (Fig 1). This selection method enabled the isolation of lines without the selection bias that was caused by the transcription levels of the *LUC* genes. However, we found differences between the results of plants and cultured cells, despite the similar experimental conditions used in the two experiments. For instance, compared with the cultured cells, plants were more prone to be transcriptionally activated by the trapping of endogenous gene/transcripts (Figs 2D and 3B), and the transcriptional strength of such activated transcription tended to be bipolarized to lower and higher transcription levels according to the insertion type (Fig 2F). How can these features of T2-plants be explained? Although transgenic cultured cells were regarded as the T1 generation, we used the T2 generation of transgenic plants in this study. Plants require a greater number of genes than do cultured cells during this one-cycle generation, because plants experience germination, development, differentiation and sexual reproduction, while the cultured cells are only in the state of vegetative propagation in a constant culture condition. Gene-insertion events might cause lethal effects on a certain population of transgenic plants by disrupting various genes that are essential for their growth over the life cycle [53]. Therefore, although we assumed that the T2-plant lines were established under a non-selective condition for *LUC* activity, the population might be distorted through a generation. Km-based selection might also affect the T2-plant population; T-DNA insertion sometimes fails to confer Km resistance and causes embryonic lethality [54,55]. In addition, under the selective condition, T-DNAs tended to be inserted in open-chromatin and hypomethylated regions [56]. Thus, Km-based selection might enrich transgenic lines in which inserts were located in the transcriptionally permissive regions where the Km-resistance genes could function sufficiently. We observed a weak insertion preference of *LUC* genes in the accessible chromatin regions (S1 Table) by utilizing a Plant Chromatin State Database [57] (http://systemsbiology.cau.edu.cn/chromstates). However, we could not evaluate any clear correlation between the transcriptional activation of promoterless *LUC* genes and the chromatin states of the corresponding WT loci, probably because the detected *LUC* population was not sufficiently large for such an analysis. Overall, the transcriptional fates of promoterless *LUC* inserts were likely to be affected by the experienced life stages and selective conditions during the establishment of transgenic plants. Hence, to grasp the extent to which inserted promoterless coding genes actually become transcribed in plants, alternative experimental strategies are needed; for example, selection-free transformation or the use of a binary vector system to introduce reporter and selection marker genes independently [58].

To date, studies of the evolutionary processes by which genetic novelty emerges were mainly led by comparative genomics [19,24,59–61]. However, because such genomics approaches are established based solely on the evolutionary winners, the resultant scenario lacks perspective from the great majority of evolutionary losers. The resolution depends on the divergence time, ranging from millions to billions of years. Conversely, our artificial evolutionary approach sheds light even on evolutionary losers within a much shorter timescale. Specifically, as the *LUC* genes used in this study are not profitable for plants, most of them would presumably become silenced or pseudogenized, while a few of them might occasionally be retained. How many generations and populations are needed to reach such endings? Our approach based on the use of plants will reveal the types of genomic/epigenomic variations that become winners/losers, thus enabling the tracing of the fates of newly activated transcripts in the population over the generations. In contrast, the cultured cells will be a useful model to investigate the molecular mechanisms underlying promoter birth, thus providing a homogeneous and simple experimental system.

It is intriguing to utilize a stress-tolerance/inducible gene as a promoterless reporter gene in our artificial evolutionary experiment. This would be a useful model to investigate how newly emerged genes adapt and evolve against exposed stress or selective environments. It is also interesting to try such experiments among different developmental phases and tissues. For example, the promoterless genes might be more prone to be transcribed in the pollen, where new genes often arise because of the transcriptionally permissive status caused by the accessible chromatin configuration [62]. Such an approach allows the investigation of gene evolution in multicellular organisms, thus providing insights into how newly emerged genes become integrated into pre-existing spatio-temporal genetic networks.

In conclusion, our artificial evolutionary experiment provided insight into the initial genetic behaviour of newly activated transcription in the plant genome. We showed that the *de novo*-activated transcription accompanying the local chromatin remodelling was inheritable. To evaluate the contribution of this phenomenon to the plant genome evolution, examination of the genetic behaviour of the *de novo* transcribed genes over an increasing number of generations with/without selective pressures will provide further clues regarding this phenomenon.

## Materials and Methods

### Plant materials and transformation

*A. thaliana* (ecotype; Col-0) plants were grown at 23°C with continuous illumination (20–50 µmol m^−2^ s^−1^). Ti-plasmid libraries containing short sequences (5’-aggcctcgacgttatcagcttacag-3’), a 12 bp random sequence (‘barcode’), a promoterless *LUC*-coding sequence, a nos-terminator and an expression cassette of a kanamycin (Km)-resistance gene between the left (LB) and right (RB) borders of the T-DNA were constructed using a modified pGreenII vector [63,64]. *Agrobacterium tumefaciens* (GV3101) cells were transformed with the Ti-plasmid libraries. *Agrobacterium*-mediated transformation of *A. thaliana* was performed according to the floral-dip method [37]. Transformed seeds were selected on Murashige and Skoog (MS) medium [1× strength of MS plant salt mixture (Nihon Pharmaceutical), 1% sucrose, 0.05% MES, 0.8% agar, pH 5.7] supplemented with 25 µg ml^−1^ of Km. The screened 386 individual Km-resistant T1 seedlings were grown at 23°C with continuous illumination (20–50 µmol m^−2^ s^−1^). The seeds of individual T2-generation plants were harvested. For the promoter-trap experiment, three seeds of individual T2-plants were stratified at 4°C in the dark for 2 days, then grown on MS medium [half-strength MS medium including vitamins (Duchefa Biochemie), 1% sucrose, 0.8% agar, pH 5.7] at 23°C with continuous illumination (40–60 µmol m^−2^ s^−1^) for 10 days. All seedlings were harvested and ground in liquid nitrogen to a fine powder, for thorough mixing. DNA and RNA were extracted using a DNeasy Plant Mini Kit (QIAGEN) and RNeasy Plant Mini Kit (QIAGEN), respectively, and subjected to the preparation of the NGS libraries.

### Determination of the *LUC* insertion sites

NGS libraries for determining *LUC* insertion loci were prepared according to the TRIP method [38,65] with modifications as follows. Genomic DNA (2.0 µg) was digested completely with DpnII, MseI or ApoI, and then purified using the QIAquick PCR purification Kit (QIAGEN). Each digested DNA (600 ng) was independently circularized with T4 DNA ligase. An aliquot of each circularized DNA was subjected to inverse PCR using primer sets that hybridize within the *LUC* ORF. Subsequently, NGS libraries were prepared by two rounds of PCR; the first round was performed to add Illumina adapters, and the second was carried out using Nextera XT index primers (Illumina). Sequencing was performed using a 301 bp paired-ended protocol on an Illumina MiSeq platform. All primers used in this study are listed in S2 Table.

For the determination of each *LUC* insertion site, NGS reads were first processed, before mapping to the genome according to a method previously described [27], with the following modifications. NGS reads were aligned to the T-DNA vector sequence (5’-tcaaggcctcgacgttatcagcttacagNNNNNNNNNNNN<u>ATGGAAGACGCCAAAAACATAAAGAAAGG CCCGGCGCCATTCTATCCTCTAGAG</u>-3’; lowercase, border sequence; N, barcode; underlined, *LUC* fraction) using Blastn (version: 2.4.0+) [66], to obtain individual flanking sequences from the *LUC* insert and barcode. The obtained flanking sequences were mapped on the TAIR10 version of the *A. thaliana* genome using bowtie [67] allowing three mismatches. Precise locus–barcode pairs were determined according to the following criteria: (1) at least two read counts; (2) the read count of the most frequent locus–barcode pair accounted for ≥60% of them, including their PCR/sequencing artefacts; and (3) exclusion from subsequent analysis of two or more distinct *LUC* inserts with the same barcode sequences. The insertion loci of T2-plants were classified according to the TAIR10 version of the genomic annotation of *A. thaliana* under the following classification: genomic regions where annotated protein-coding genes were defined as ‘Genic’ regions, whereas the remainder of the genome was classified as ‘Intergenic’. The insertion strand of *LUC* genes was considered.

### Determination of the relative transcription level of *LUC* genes

NGS libraries for determining *LUC* transcription level were prepared according to the TRIP method [38,65] with modifications as follows. RNA (5.0 µg) was subjected to reverse transcription using an oligo dT_15_ primer and SuperScript III Reverse Transcriptase (Thermo Fisher Scientific). An NGS library (termed RNA library, hereafter) was prepared by amplification of the barcode region of *LUC*-cDNA using primer sets with an Illumina adapter extension, followed by the tailed-PCR using Nextera XT index primers. From an aliquot of DNA used in the *LUC* insertion site determination, barcode regions were amplified and an NGS library (termed DNA library, hereafter) was prepared according to the method described above. Sequencing was performed on the Illumina MiSeq under a 76 bp paired-ended protocol.

To obtain an indicator of the molecular abundances of each *LUC*-mRNA per transgenic cells, barcode sequences were extracted from the sequencing reads and counted. Barcodes with a read number ≤5 in the DNA library were omitted from further analysis. In the RNA library, barcodes with a read number ≤5 were set as zero. For DNA and RNA libraries, the read number of each barcode was normalized to the total sequencing reads of the corresponding library. The relative transcription level of each *LUC* gene was calculated as follows: the RNA read number of each barcode was divided by the corresponding DNA read number and multiplied by 10,000. Subsequently, individual *LUC* loci and transcription levels were associated based on their barcode sequences.

### Validation of *LUC* insertion loci and barcode sequences

Randomly chosen T2-plants were stratified at 4°C in the dark for 2 days, then grown on MS medium supplemented with 25 µg ml^−1^ of Km at 23°C with continuous illumination (20–30 µmol m^−2^ s^−1^) for 10 days. Km-resistant seedlings were harvested and subjected to DNA extraction. Four types of PCR were performed to amplify the barcode region, the RB–genome junction, the LB–genome junction and the T-DNA insert, respectively. The PCR products obtained were then analysed by agarose gel electrophoresis and Sanger sequencing, for validation of the insertion locus and barcode sequence, respectively.

### Comparison with WT transcriptome data

RNA-seq data of WT *A. thaliana* (col-0) plants were retrieved from the *NCBI Short-Read Archive* under accessions SRR6388204, SRR6388205 and SRR770510. The sequencing reads were subjected to adapter trimming and quality trimming, followed by mapping to the *A. thaliana* genome (TAIR10) using STAR (v2.5.3) [68] with the following parameters: *STAR – alignIntronMax 6000 –outSAMstrandField intronMotif –two passMode Basic*. Transcribed regions and their transcription levels (in fragments per kilobase of exon per million reads mapped (FPKM)) were analysed using StringTie (v2.1.4) [69]. Subsequently, the transcription level of each T2-plant was compared with the FPKM of the inherent transcribed region in the WT genome. In the case of inherent transcripts with multiple isoforms, each FPKM was summed up.

### Chromatin immunoprecipitation (ChIP) and MBD immunoprecipitation (MBDIP) analysis

The T2:161 and T2:205 lines were stratified at 4°C in the dark for 3 days, then grown on MS medium [half-strength MS medium including vitamins (Duchefa Biochemie), 1% sucrose, 0.8% agar, pH 5.7] supplemented with 15 μg ml^−1^ of Km at 23°C with continuous illumination (20–30 µmol m^−2^ s^−1^) for 8 days. Km-resistant seedlings were harvested and subjected to ChIP and MBDIP analysis. For control experiments, transgenic *A. thaliana* harbouring an expression cassette of the Km-resistance gene without the *LUC* reporter gene (termed WT in Figs 4D and E, and 5B) were prepared and grown under the same condition as that used for T2-plants. ChIP and MBDIP were performed according to a method previously described [27], with the following modifications. For the ChIP assay, ∽10 ng of solubilized chromatin (median, 200 bp) and antibodies (2.4 µg of an anti-H2A.Z antibody [70] and 2.0 μg of an anti-H3K36me3 antibody (Abcam: ab9050), respectively) were used for each experiment. For the MBDIP assay, the methylated DNA fraction (mC) was collected from 1.0 μg of sheared DNA (median, 200 bp) using an EpiXplore Methylated DNA Enrichment Kit (Clontech) according to the manufacturer’s instructions. Successful enrichment of ChIPed DNA and mC was validated by qPCR in the control sites (S2 Table) according to Deal *et al*. [71] for H2A.Z, to Yang *et al*. [72] for H3K36me3 and to Erdmann et al. [73] for mC. In both T2-plants and WT, relative enrichments of H2A.Z, H3K36me3 and mC around the *LUC* insertion loci were calculated based on the enrichment of the control sites, which was set as 100%.

### Expression and TSS analysis

The T2:161 and T2:205 lines were grown and harvested under the same condition as that used for the ChIP experiments. Total RNA was isolated using an RNeasy Plant Mini Kit followed by DNase I treatment. For expression analysis, cDNA was synthesized from 5.0 µg of the total RNA using an oligo dT_20_ primer and SuperScript III Reverse Transcriptase. The transcription level of the *LUC* gene was normalized to that of the ubiquitin gene (*UBQ10*: AT4G05320).

*LUC*-TSS was analysed according to a published method [52,74], with the following modifications. Specifically, polyadenylated RNA was extracted using a Dynabeads mRNA Purification Kit (Invitrogen) according to the manufacturer’s protocol. Polyadenylated RNA (1.0 µg) was used for reverse-transcription and template-switching reactions. During these reactions, SgfI sites were added at both ends of the full-length cDNA by the primer used for reverse transcription and the template-switching oligo. The full-length cDNAs obtained were then digested completely by SgfI. Subsequently, digested cDNAs were circularized and subjected to inverse PCR to specifically amplify *LUC*-containing cDNAs. The resulting nested PCR products were analysed by Sanger sequencing.

## Acknowledgements

We thank Moyuru Shirasu for his help in maintaining the transgenic *A. thaliana* plants.

## Supporting information

**S1 Fig**. Validation of *LUC* insertion loci.

**S2 Fig**. Expression analysis of T2-plants.

**S3 Fig**. Sequence alignment of TSS of T2:161 line.

**S1 Table**. List of the *LUC* lines.

**S2 Table**. Primer list.

